# A Dual Function Antibody Conjugate Enabled Photoimmunotherapy Complements Fluorescence and Photoacoustic Imaging of Head and Neck Cancer Spheroids

**DOI:** 10.1101/2023.01.30.526194

**Authors:** Mohammad A. Saad, Stacey Grimaldo-Garcia, Allison Sweeney, Srivalleesha Mallidi, Tayyaba Hasan

## Abstract

Several molecular-targeted imaging and therapeutic agents are in clinical trials for image-guided surgery and photoimmunotherapy (PIT) of head and neck cancers. In this context, we have previously reported the development, characterization, and specificity of a dual function antibody conjugate (DFAC) for multi-modal imaging and photoimmunotherapy (PIT) of EGFR over-expressing cancer cells. The DFAC reported previously and used in the present study, comprises of an EGFR targeted antibody – Cetuximab conjugated to Benzoporphyrin derivative (BPD) for fluorescence imaging and PIT, and a Si-centered naphthalocyanine dye for photoacoustic imaging. We report here the evaluation and performance of DFAC in detecting microscopic cancer spheroids by fluorescence and photoacoustic imaging along with their treatment by PIT. We demonstrate that while fluorescence imaging can detect spheroids with volumes greater than 0.049 mm^3^, photoacoustic imaging-based detection was possible even for the smallest spheroids (0.01 mm^3^), developed in the study. When subjected to PIT, the spheroids showed a dose-dependent response with smaller spheroids (0.01 and 0.018 mm^3^) showing a complete response with no recurrence when treated with 100 J/cm^2^. Together our results demonstrate the complementary imaging and treatment capacity of DFAC. This potentially enables fluorescence imaging to assess tumor presence on a macroscopic scale followed by photoacoustic imaging for delineating tumor margins guiding surgical resection and elimination of any residual microscopic disease by PIT, in a single intra-operative setting.

## Introduction

Antibody-based targeted diagnostic and therapeutic agents have been demonstrated for their utility in the treatment of several malignant disorders including head and neck cancers [1–3]. Antibody-based targeted photoactive agents are currently being employed in head and neck cancers for tumor margin assessment during surgical resection and photoimmunotherapy (NCT02422979, NCT03769506) [4–7]. Attaining disease free surgical margins while minimizing resection of healthy tissue is a major challenge in tumor resection surgeries for head and neck cancers. Precise identification of tumor margins is important in minimizing healthy tissue loss, and preserving vital structures, function, and aesthetics to improve patient compliance. As recurrence rates in head and neck cancers decrease with every extra millimeter of healthy tissue resected [8], it is important to assess these molecular-targeted probes for their ability in accurately delineating tumor margins and treating tumors of microscopic dimensions. In this context, while fluorescence imaging provides a macroscopic overview to identify extent of tumor invasion and lymph node metastasis, complementing it with photoacoustic imaging, as proposed in the present study using DFAC, can help in delineation of tumor margins in three dimensions.

Recent advances in tumor-targeted imaging have seen several antibody fluorophore conjugates being tested in clinical trials for guiding resection surgeries [9, 10]. In this context, the use of photosensitizers provides an advantage where the photoactive molecule can be used for fluorescence imaging and treatment of the target lesions through photodynamic therapy. This has been shown in several studies from our lab and others [2, 11, 12]. Although photoimmunotherapy (PIT) was developed three decades ago [13], it has recently gained popularity with clinical approval in Japan for head and neck cancers. Ongoing phase III clinical trials are evaluating the efficacy of PIT for head and neck cancers in USA [4–6, 14]. The efficacy of photodynamic therapy (PDT), however, largely depends on tumor size and oxygenation. PDT of larger or hypoxic tumors results in worse therapeutic outcomes, whereas smaller or more oxygenated tumors respond better [15–17].

We have previously reported the synthesis and characterization of a dual function antibody conjugate (DFAC) for multi-modal imaging (photoacoustic and fluorescence imaging) and PIT of EGFR over-expressing tumor cells [1, 18]. While fluorescence imaging provides sensitivity, the tumor depth profiling provided by fluorescence imaging is low, a deficiency adequately complemented by photoacoustic imaging. The ability to irradiate the tumor bed post-surgery carries the potential of eliminating microscopic disease thus minimizing recurrence. In this study, we evaluate the performance of the DFAC developed using Cetuximab (an EGFR targeted antibody) conjugated to Benzoporphyrin derivative (BPD) for fluorescence imaging and PIT, and a Si-centered naphthalocyanine (SiNC(OH)) [19] dye for photoacoustic imaging in microscopic head and neck cancer spheroids, in vitro. We also propose a framework for executing multi-modal imaging followed by PIT for precise delineation of tumor margins and elimination of residual disease.

## Materials and Methods

### Dual function antibody conjugate (DFAC) synthesis

Anti-EGFR antibody: Cetuximab (Erbitux^®^, Eli Lilly and Co., Indianapolis, IN, USA) was filtered (by passing through a 0.22 μm syringe filter) under sterile conditions. The concentration of the antibody was estimated using a Pierce™ BCA Protein Assay Kit (ThermoFisher Scientific, Waltham, MA, USA). Approximately 4 mg of antibody (2 mL of antibody) was obtained in a scintillation vial and 0.2 mL of 4 mg/ml Methoxy PEG Succinimidyl Carbonate, (MW 10000) (JenKem Technology, Plano, TX, USA), dissolved in DMSO, was added drop wise under stirring. The reaction was stirred over night at room temperature. SiNC(OH)-NHS and BPD-NHS (at 4 times molar excess), prepared as described in our previous studies [1, 19, 20], dissolved in DMSO, were added dropwise to the PEGylated antibody and the reaction was allowed to proceed for 4 h. The total DMSO content in the reaction mix was never allowed to exceed 40%. The reaction mix was centrifuged for 10 min at 15000 *x* g to remove any insoluble aggregates formed during the reaction. The supernatant containing the crude DFAC was passed through a Zeba™ Spin Desalting Column (7K MWCO, 10 mL) (ThermoFisher Scientific, Waltham, MA, USA) pre-equilibrated with 30% DMSO. The eluent obtained after passing through the gel filtration column was then subjected to buffer exchange and concentrated using a 30 kDa NMWL Amicon^®^ Ultra-15 Centrifugal Filter Unit (MilliporeSigma, Burlington, MA, USA). The DMSO was approximately 5% in the final DFAC product. DFACs were stored at 4 °C and remained stable for several months.

### Estimation of BPD and SiNC(OH) molar loading ratios in DFACs

Absorption spectra of DFAC was recorded, at appropriate dilutions in DMSO, using an Evolution™ 300 UV-Vis Spectrophotometer (ThermoFisher Scientific, Waltham, MA, USA). Absorbance at 687 nm and 872.6 nm were recorded and the respective concentrations of BPD (34,895 M^−1^cm^−1^) and SiNC(OH) (277,095 M^−1^cm^−1^) were calculated using their extinction coefficients in DMSO. The concentration of the Cetuximab antibody was calculated using a Pierce™ BCA Protein Assay Kit (ThermoFisher Scientific, Waltham, MA, USA). The loading ratios and conjugation efficiencies of the two dyes were subsequently calculated using the obtained concentrations of the dyes and protein (antibody), respectively.

### Oral squamous cell carcinoma line (SCC-4) culture and generation of EGFP expressing SCC4 cells

Human oral squamous cell carcinoma line SCC4 was obtained from American Type Culture Collection (ATCC CRL-1624™, Manassas, VA, USA). All cells were cultured in DMEM:F12 Medium (SH30271.01, Cytiva, Marlborough, MA, USA) supplemented with an antibiotic mixture containing Penicillin (100 I.U/mL) and streptomycin (100 μg/mL) (30-001-Cl, Corning, Corning, NY, USA), 10% heat-inactivated fetal bovine serum (FBS) (SH30071.03HI, Hyclone™, Marlborough, MA, USA) and 400 ng/mL hydrocortisone (MilliporeSigma, Burlington, MA, USA). The transgenic EGFP expressing variant of SCC4 cell line was generated by a previously established protocol [2]. Briefly, third-generation EGFP transfer plasmid pHAGE-CMV-EGFP-W (EvNO00061634, Harvard Plasmid Repository) was mixed with viral envelope plasmid pMD2.G (12259, Addgene, Watertown, MA) and viral packaging plasmid psPAX2 (12260, Addgene, Watertown, MA) in a molar ratio of 2:1:1 and transfected into the Lenti-X™ 293T packaging cell line (632180, TaKaRa Bio Inc., Kusatsu, Shiga, Japan) using Xfect™ Transfection Reagent (631317, TaKaRa Bio Inc., Kusatsu, Shiga, Japan) following manufacturer’s instruction. The viral supernatant was collected 48 h post-transfection, filtered using a 0.45 μm Puradisc 25 mm polyethersulfone Syringe Filter (6780-2504, GE Healthcare Life Sciences, Whatman™, Chicago, IL, USA) to remove cellular debris, and concentrated using a Lenti-X™ Concentrator (631231, TaKaRa Bio Inc., Kusatsu, Shiga, Japan). SCC4 cells were then transduced with viral suspension at different multiplicity of infection (MOI) in the presence of 6 μg/mL hexadimethrine bromide (polybrene, MilliporeSigma, Burlington, MA, USA). Cells were sorted for EGFP fluorescence using a BD FACSAria™ Cell Sorter (BD Biosciences, San Jose, CA, USA), after 48 h of transduction. The sorted cells were expanded and cryopreserved in fetal bovine serum (Gibco™, ThermoFisher Scientific, Waltham, MA, USA) with 8% DMSO. All cell lines used in this study tested negative for Mycoplasma contamination via MycoAlert™ PLUS Mycoplasma Detection Kit (Lonza, Basel, Switzerland).

### Preparation of tumor spheroid encapsulating tissue phantom gelatin molds for fluorescence and photoacoustic imaging

SCC4-EGFP spheroids were grown as described earlier and tumor spheroid containing phantoms were prepared according to previously described protocols [1, 21]. Briefly, 24 h post cell-seeding, the spheroids were treated with different concentrations of DFAC (0.25 μM, 1 μM and 2.5 μM; SiNC(OH) equivalent). After 24 h, the spheroids were washed with PBS thrice, and placed on top of a gelatin bed (prepared from 10% gelatin) in a well of a 96-well plate. After a few minutes, 10% gelatin was carefully pipetted on top of the spheroid and allowed to cool/solidify and imaged within 24 h.

### Imaging of tumor spheroid embedded tissue phantoms

The tumor spheroid embedded tissue phantoms were imaged for their fluorescence intensity, exploiting the Q-band of Verteporfin, using an IVIS^®^Lumina Series III In Vivo Imaging System (Perkin Elmer, Waltham, MA). Excitation and emission filters of 660 nm and 710 nm, respectively. Quantification of fluorescence intensities were performed using the Living Image^®^ software provided by the manufacturer. Background fluorescence was calculated by imaging untreated spheroids. Tumor to background ratios were calculated by dividing the fluorescence signal intensity of the treated spheroids with the fluorescence signal intensities of the untreated spheroids of the same size. Photoacoustic imaging (PAI) was performed by submerging the 96 well plate in water and placing the PA transducer (LZ220) in close proximity to the spheroid containing phantoms. Imaging parameters - laser power, photoacoustic gain, persistence, and frame rate were kept constant across different measurements. Image analysis was performed using the built-in software provided by Visualsonics, Fujifilm (ON, Canada). Ultrasound and photoacoustic raw data for various spheroids were exported to MATLAB. A custom written GUI was used to segment the nodule in the ultrasound image. The segmentation mask was then overlaid on PA image obtained at various wavelengths. Owing to the small size of the nodules, the ultrasound, and photoacoustic images from various conditions (seeding density and concentration of the dye) were scaled 5 times (cubic interpolation) for display purposes. All images are displayed on the same dynamic range.

### Evaluation of light dose-dependent toxicity in in vitro tumor spheroids

For determining the phototoxicity of DFAC on 3D SCC4-EGFP spheroids, spheroids were cultured by seeding cells in Corning Costar Ultra-Low attachment 96 well plates in different densities (2500, 5000, 10000 and 25000 cells in 100 μL) to attain tumor volumes of different sizes. The spheroids were then treated with DFAC (0.25 μM BPD equivalent) for 24 h followed by washing with culture media thrice. The spheroids were then irradiated with a 690 nm light, administered vertically through the plate bottom via a diode laser source at 150 mW/cm^2^ irradiance as described previously [1]. Total fluence was varied from 10 J/cm^2^ to 100 J/cm^2^ by varying the irradiation time. Following irradiation, culture plates were incubated for 3 days after which they were imaged with an Operetta CLS high-content analysis system (Perkin Elmer, LIVE configuration) with the incubation chamber set to 37 °C and 5% CO_2_. EGFP (ex/em: 475/500-570 nm) fluorescence was collected via LED excitation (at 475 and 550 nm, respectively) and a 10X, 0.3 NA air objective in a z-stack format. For quantification, 2D images were generated via maximum intensity projection (MIP) to analyze in-focus light from the full z-stack. Images were analyzed via custom Image J script that applies the following analysis scheme to each well from each fluorescence channel to generate fluorescence volumetric data. Images were converted to an 8-bit format and subjected to background subtraction using a rolling ball algorithm with pixel size of 25. Equal brightness/contrast enhancement was performed to increase histogram binning. The mean fluorescence of the corrected histograms from the treated spheroids was computed and compared to the control untreated spheroids. Relative viabilities were calculated by dividing fluorescence intensity of SCC4-EGFP spheroids from the treated spheroids by the intensity of the respective untreated control SCC4-EGFP spheroids (from the same day) developed from the same SCC4-EGFP cell seeding densities.

### Tumor recurrence monitoring

For monitoring tumor recurrence, tumor spheroids were imaged longitudinally for 5 weeks following treatment and the intensity of the treated/untreated control spheroids were calculated as mentioned in the previous section. Spheroids were considered to be recurring if they showed a relative viability of 0.3 or more. Recurrence proportion was calculated by counting the number of spheroids with a relative viability of ≥ 0.3 and dividing it by the total number of spheroids.

## Results

### Synthesis and characterization of dual function antibody conjugates

The dual function antibody conjugate (DFAC) synthesis was carried out in 40% DMSO as described previously [1]. While 40% DMSO may affect the structure and function of most proteins, previous studies from our lab have established this to be an optimal concentration to maintain antibody function and achieve appropriate loading ratios for hydrophobic dyes [2, 11, 20, 22]. PEGylation of Cetuximab was performed to minimize aggregation and enhance solubility of the DFAC (**Figure 1A**) [20, 22, 23]. Through this procedure molar ratios of 2.65 ± 0.42 and 2.96 ± 0.58, per mole of antibody, were consistently obtained for BPD and SiNC(OH), respectively. While BPD was used for its well-established fluorescence and photosensitization properties, a naphthalocyanine derivative SiNC(OH) was preferred as a photoacoustic dye,[19] over the clinically used dyes indocyanine green (ICG) [24], methylene blue (MB) [25], and IRDye800 [26] due to its near infra-red absorption profile which does not overlap with the absorption spectrum of oxygenated/deoxygenated hemoglobin. Moreover, naphthalocyanine dyes have high extinction coefficients and a high non-radiative quantum yield [19, 27, 28], as compared to the above-mentioned dyes, and hence can be used for improving photoacoustic contrast.

**Figure 1:**
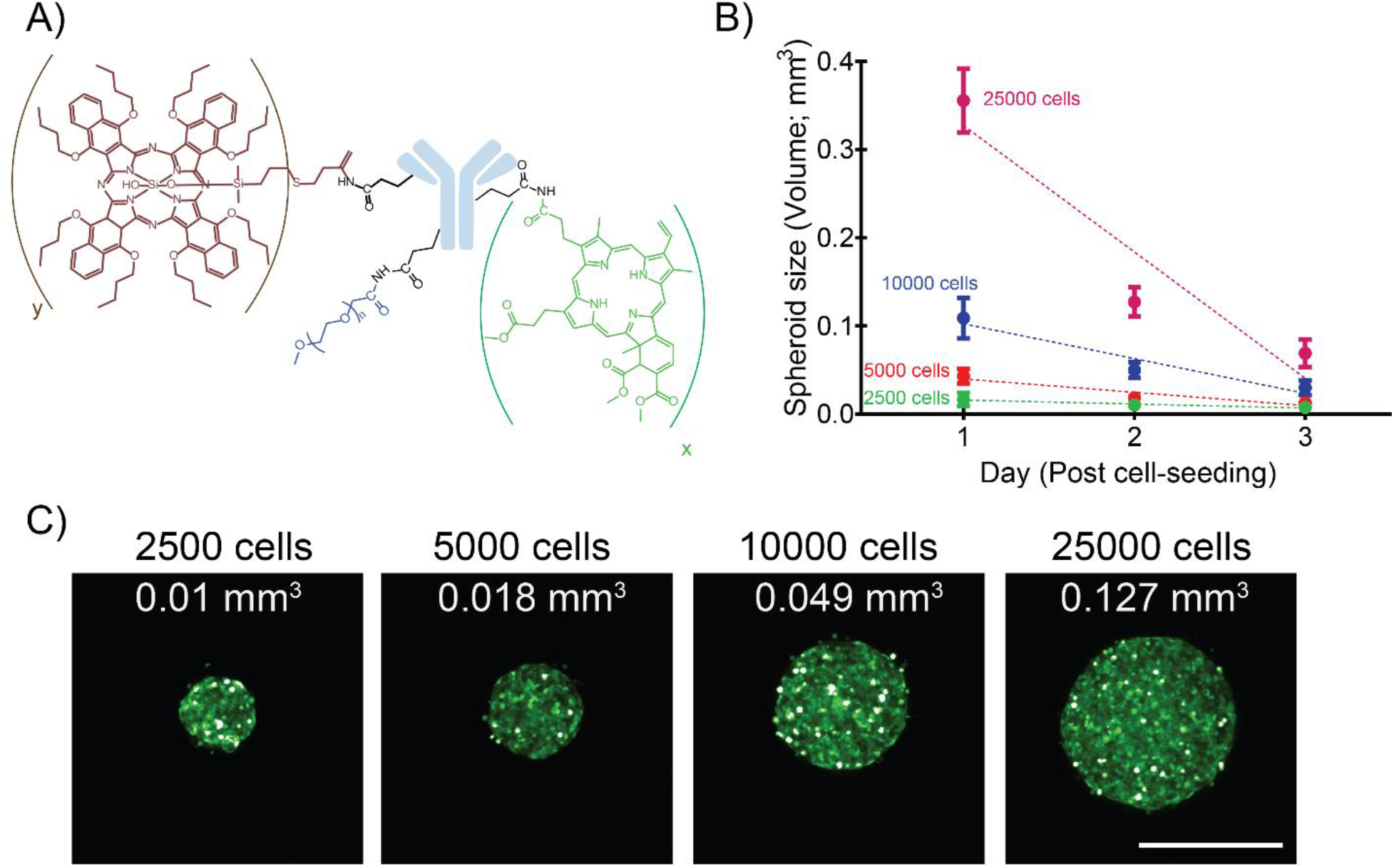
Characterization of SCC4-EGFP spheroids with different volumes. **(A)** Structure of the dual function antibody conjugate. The number of BPD and SiNC(OH) molecules per antibody molecule are denoted by “x” and “y” respectively. **(B)** Graph representing the change in SCC4-EGFP spheroid volume on different days, post cell-seeding. **(C)** Representative images of SCC4-EGFP 3D spheroids formed with different cell numbers (2500, 5000, 10000 and 25000 cells). Cell numbers are mentioned at the top while spheroid volumes are mentioned within the images. (Scale bar = 500 μm).

### Characterization of tumor spheroids

To recapitulate microscopic tumors of different sizes, SCC4 spheroids were developed by seeding different numbers of cells (2500, 5000, 10000 and 25000) per well of Corning Costar Ultra-Low attachment 96 well plates. The SCC4 cells were transduced with an EGFP lentiviral expression vector to enable fluorescence-based longitudinal monitoring of the spheroids and assessment of viability [2, 29]. Spheroid volumes were calculated using the formula (V = 0.5 * Length * Width^2^), as reported previously [30, 31]. As evident from **Figure 1B**, spheroid volume increased with increasing cell numbers. While the volumes for spheroids formed with lower cell numbers (2500 and 5000) did not change significantly over time, volumes of spheroids formed with 10000 and 25000 cells decreased from 0.11 ± 0.02 mm^3^ and 0.36 ± 0.04 mm^3^ on day 1 to 0.03 ± 0.01 mm^3^ and 0.07 ± 0.02 mm^3^ by day 3, respectively (**Figure 1B**). **Figure 1C** provides representative images of the spheroids (on day 2 post seeding). The morphology of the spheroids was consistent with that reported previously for SCC4 [32] and the significant decrease in size, with time, for spheroids formed with higher cell numbers can be attributed to a progressive slow death phenotype [33]. This is possibly due to the hypoxic core formed in these spheroids, as has been reportedly previously for several oral cancer cell lines, which results in the formation of a necrotic zone in the spheroid center [33].

### Fluorescence Imaging of tumor spheroids

Fluorescence imaging of the spheroids was performed according to the scheme provided in **Figure 2A**. After treatment of spheroids with DFAC, with the indicated concentrations for 24 h, the spheroids were washed with PBS and tissue phantoms were prepared followed by fluorescence imaging through an IVIS^®^ Spectrum in vivo imaging system. As shown in **Figure 2B**, fluorescence signals were observed from spheroids from nodules with volumes >0.049 mm^3^ treated with a DFAC concentration of 2.5 μM SiNC(OH) equivalent. Spheroids with volumes of 0.127 mm^3^ also showed a fluorescence signal when treated with 1 μM DFAC (SiNC(OH) equivalent). **Figure 2C** shows the tumor spheroid fluorescence to background ratio for the different spheroids treated with increasing concentrations of DFAC. Treatment with 2.5 μM DFAC ((SiNC(OH) equivalent) resulted in a significantly higher fluorescence intensity for spheroids with volumes >0.018 mm^3^. The low fluorescence intensity from the spheroids with smaller volumes (lower cell numbers) could be due to the lower amount of the fluorophore (BPD) accumulated in these spheroids, the low extinction coefficient of BPD (34,895 M^−1^cm^−1^), and the lower fluorescence quantum yield of BPD (~0.05) [34]. These results provide us with a general idea of the lowest tumor volumes that can be detected with different concentrations of DFAC used.

**Figure 2:**
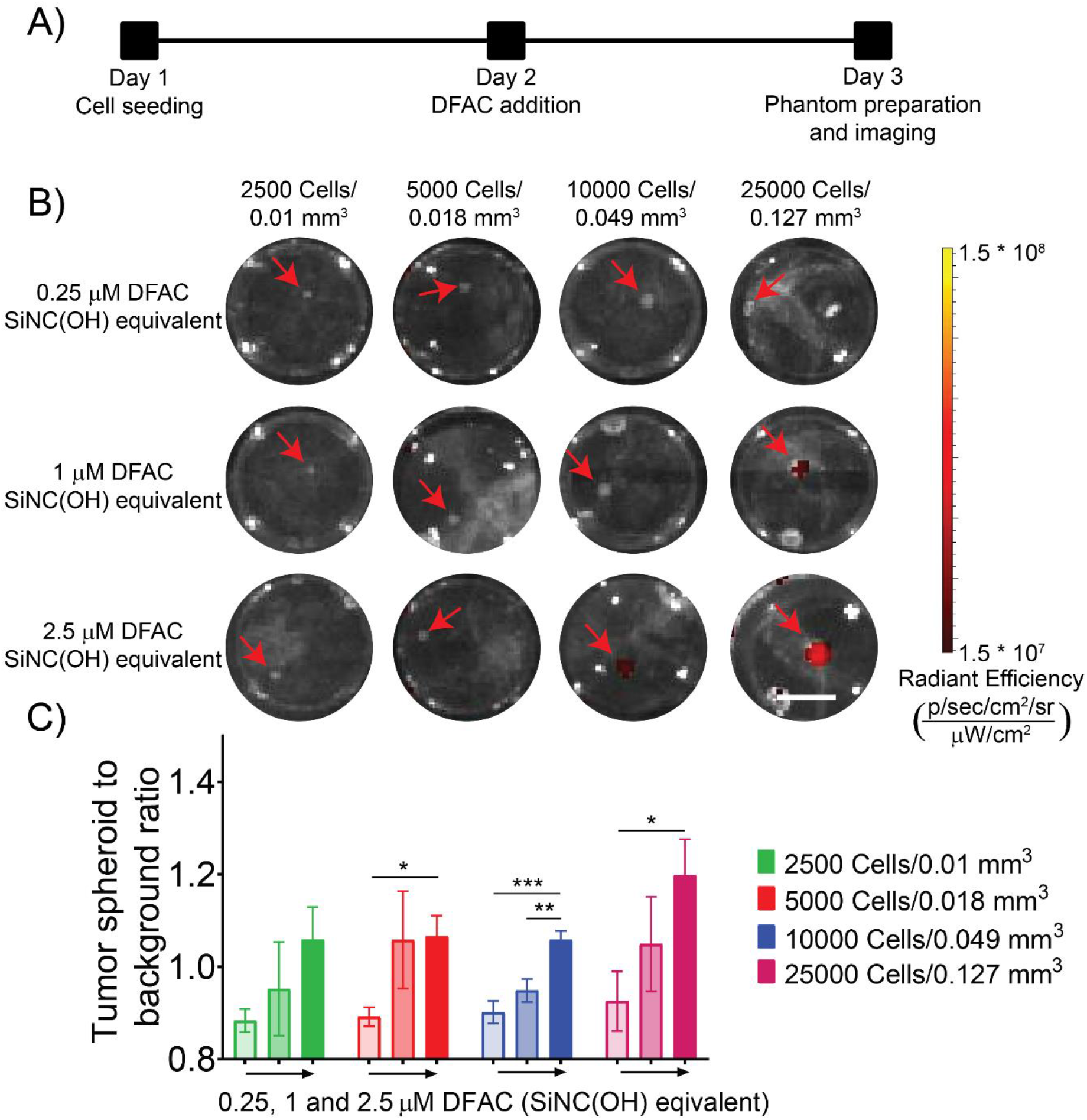
Fluorescence imaging of DFAC-treated SCC4 spheroids: **(A)** Experimental scheme of SCC4-EGFP 3D spheroid treatment with DFAC and imaging. **(B)** IVIS fluorescence imaging of DFAC treated SCC4 spheroids prepared as gelatin embedded tissue mimicking phantoms in a 96-well plate (Scale bar = 2.5 mm). Cell numbers and volumes are mentioned at the top while DFAC concentrations (SiNC(OH) equivalent) are mentioned on the left. **(C)** Quantification of fluorescence signals from the IVIS images represented as tumor spheroid to background ratio. Background fluorescence signals were obtained by quantifying fluorescence signal from control untreated wells. Fluorescence signal from spheroids increased with increase in spheroid size and DFAC concentrations. Data are presented as mean ± SD (n ≥ 3), analyzed using one-way ANOVA with Tukey ‘s test for post hoc analysis. p-values < 0.05 were considered to be significant and are indicated by asterisks as follows: ^ns^p > 0.05, *p < 0.05, **p < 0.01, ***p < 0.001 and ****p < 0.0001.

### Photoacoustic imaging of tumor spheroids

We next performed photoacoustic imaging of the cell phantoms. As shown in **Figure 3A**, the photoacoustic signal from the SCC4 spheroids treated with DFAC was detectable even for the smallest spheroids with a volume of 0.01 mm^3^. Spectral photoacoustic scanning was performed to distinguish spheroids from background noise originating from air bubbles in the phantom. In general, the PA signal intensity increased with increase in the DFAC concentration for spheroids of all sizes. The PA signal intensity was comparable between all spheroids treated with the same DFAC concentration (**Supplementary Figure 1**). However, as shown in **Figure 3B** smaller spheroids showed a higher PA signal intensity per unit volume (mm^3^) as compared to the larger spheroids which showed relatively lower PA signal intensity per unit volume (mm^3^). While the PA signal intensity per mm^3^ for spheroids with tumor volumes of 0.01 mm^3^, 0.018 mm^3^ and 0.049 mm^3^ did not differ significantly, the PA signal intensity for tumor volume 0.127 mm^3^ was significantly lower (*p* value 0.0022) (**Figure 3B**). Combined with the fluorescence imaging where spheroids with larger size showed a higher tumor to background fluorescence signal, the complementary PA signals highlight the utility of DFACs in identifying tumors of different sizes.

**Figure 3:**
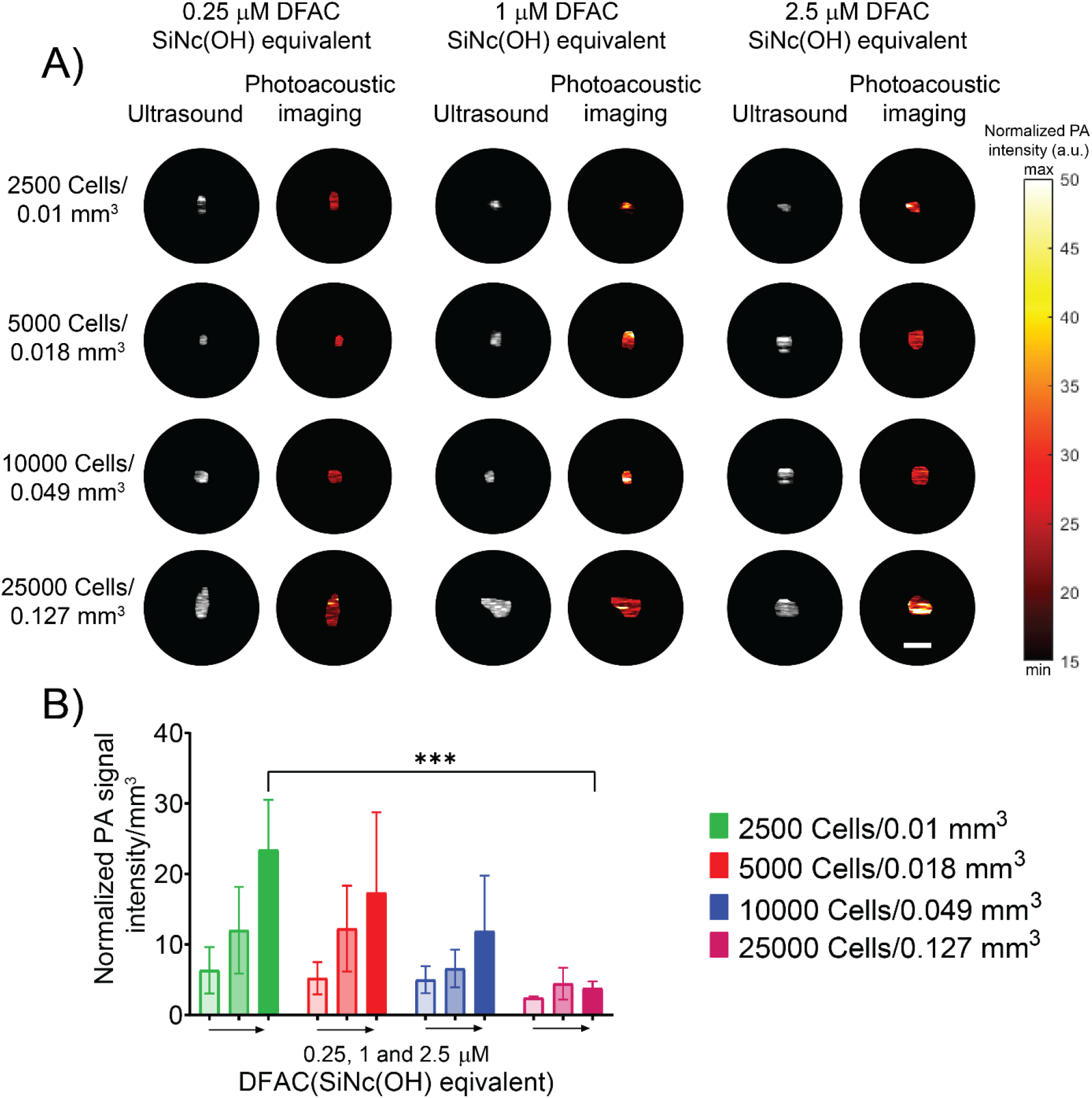
Photoacoustic imaging of DFAC treated SCC4 spheroids: **(A)** Photoacoustic imaging of DFAC treated SCC4 spheroids prepared as gelatin embedded tissue mimicking phantoms in a 96-well plate (Scale bar = 2.5 mm). Cell numbers and volumes are mentioned on the left while DFAC concentrations (SiNC(OH) equivalent) are mentioned on the top. **(B)** Quantification of photoacoustic signals from the images represented as normalized photoacoustic signal intensity per mm^3^. Spheroid volumes were obtained by performing a 3D photoacoustic scan of the spheroids. Photoacoustic signal from spheroids decreased with increase in spheroid size and the difference was more pronounced for spheroids treated with a high (2.5 μM) DFAC concentrations (SiNC(OH) equivalent). Data are presented as mean ± SD (n ≥ 3), analyzed using one-way Brown-Forsythe and Welch ANOVA test with post hoc analysis. p-values < 0.05 were considered to be significant and are indicated by asterisks as follows: ^ns^p > 0.05, *p < 0.05, **p < 0.01, ***p < 0.001 and ****p < 0.0001.

To understand the relation between DFAC uptake in the spheroids and its detection by either fluorescence or PA imaging, we quantified the cellular DFAC uptake (as monitored by BPD fluorescence) in the spheroids and determined its correlation with the fluorescence and PA signal intensity. As shown in **Figure 4**, while the total DFAC amount (determined as the product of median BPD fluorescence (a.u) and total cell number on day 2) in the spheroids increased with an increase in cell number/spheroid size (**Figure 4A**), the median BPD fluorescence decreased with increase in nodule size. This suggests that although the DFAC concentration per cell in smaller spheroids is higher it may still be below the limit of detection (LOD) of the fluorescence imaging system used in the study (IVIS) (**Figure 4B**). However, the low DFAC content in the smaller spheroids was still detectable through PA imaging and the PA signal intensity/mm^3^ for these smaller spheroids was significantly higher as compared to the larger spheroids, highlighting the complementary nature of the probe.

**Figure 4:**
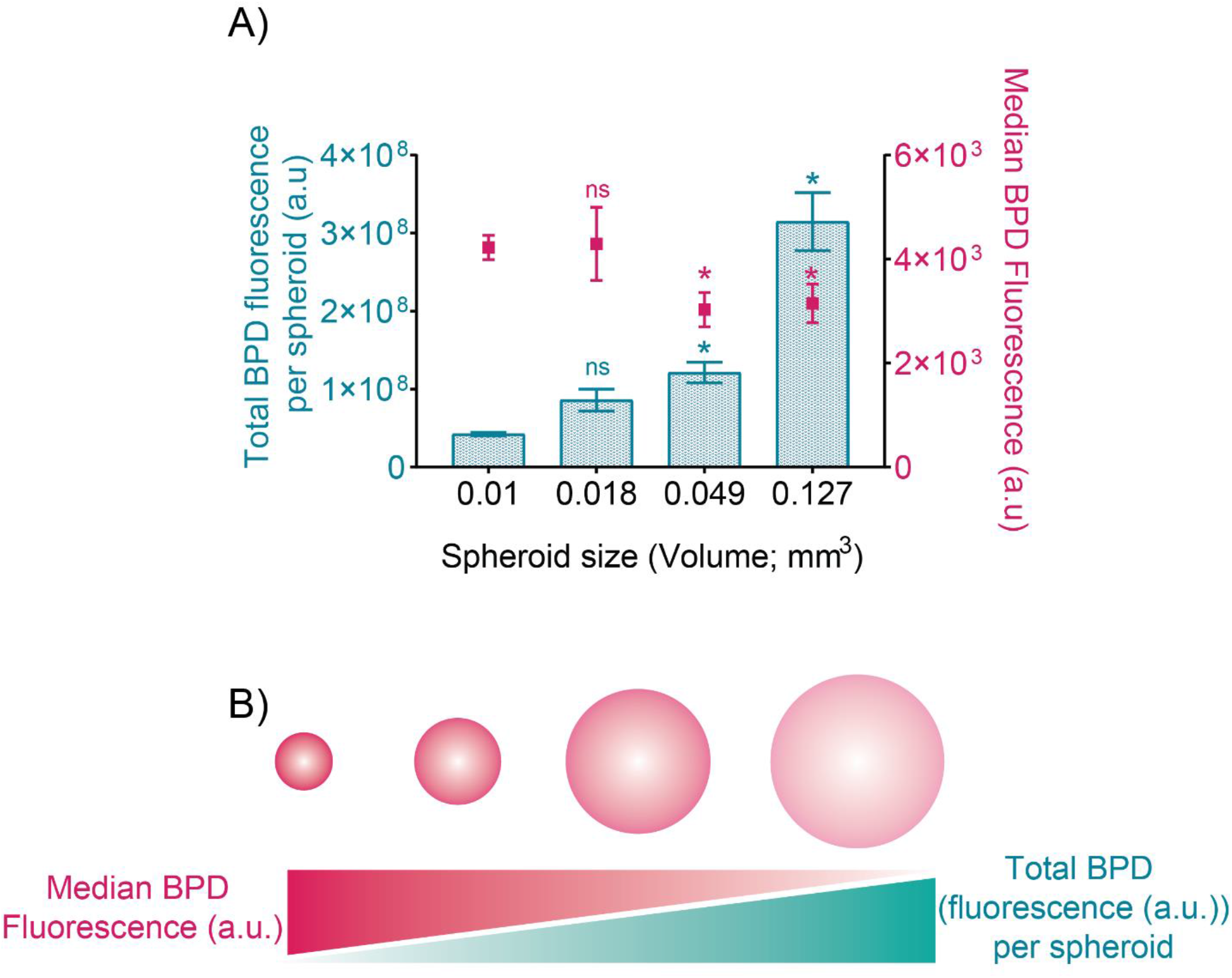
DFAC uptake in spheroids: **(A)** Total BPD fluorescence per nodule (a.u.) (cyan bar graphs) and median BPD fluorescence (a.u.)/cell in the spheroids plotted as function of spheroid size. While the total BPD fluorescence per nodule was higher for the spheroids with larger size, the median BPD fluorescence (a.u.) per cell was low. **(B)** Figurative representation of BPD distribution and BPD density in the spheroids with increasing spheroid size. As depicted, BPD density in the smaller spheroids is higher while the cumulative BPD content is higher in the larger spheroids. Data are presented as mean ± SD (n ≥ 3), analyzed using one-way Brown-Forsythe and Welch ANOVA test with post hoc analysis. p-values < 0.05 were considered to be significant and are indicated by asterisks (as compared to the 0.01 mm^3^ spheroids) as follows: ^ns^p > 0.05, *p < 0.05, **p < 0.01, ***p < 0.001 and ****p < 0.0001.

Next, we determined the correlation of PA signal intensity/mm^3^ and tumor spheroid fluorescence to background ratio with the total DFAC content per spheroid and median DFAC (BPD) fluorescence, as determined by flow cytometry. **Figure 5** shows the correlation curves of these parameters. The tumor to background fluorescence signal positively correlated with the total DFAC content of the spheroid (**Figure 5A**), while it negatively correlated with the median DFAC (BPD) fluorescence (**Figure 5C**). In contrast, the PA signal intensity/mm^3^ negatively correlated with median DFAC (BPD) fluorescence and positively correlated with DFAC content (**Figure 5D**).

**Figure 5:**
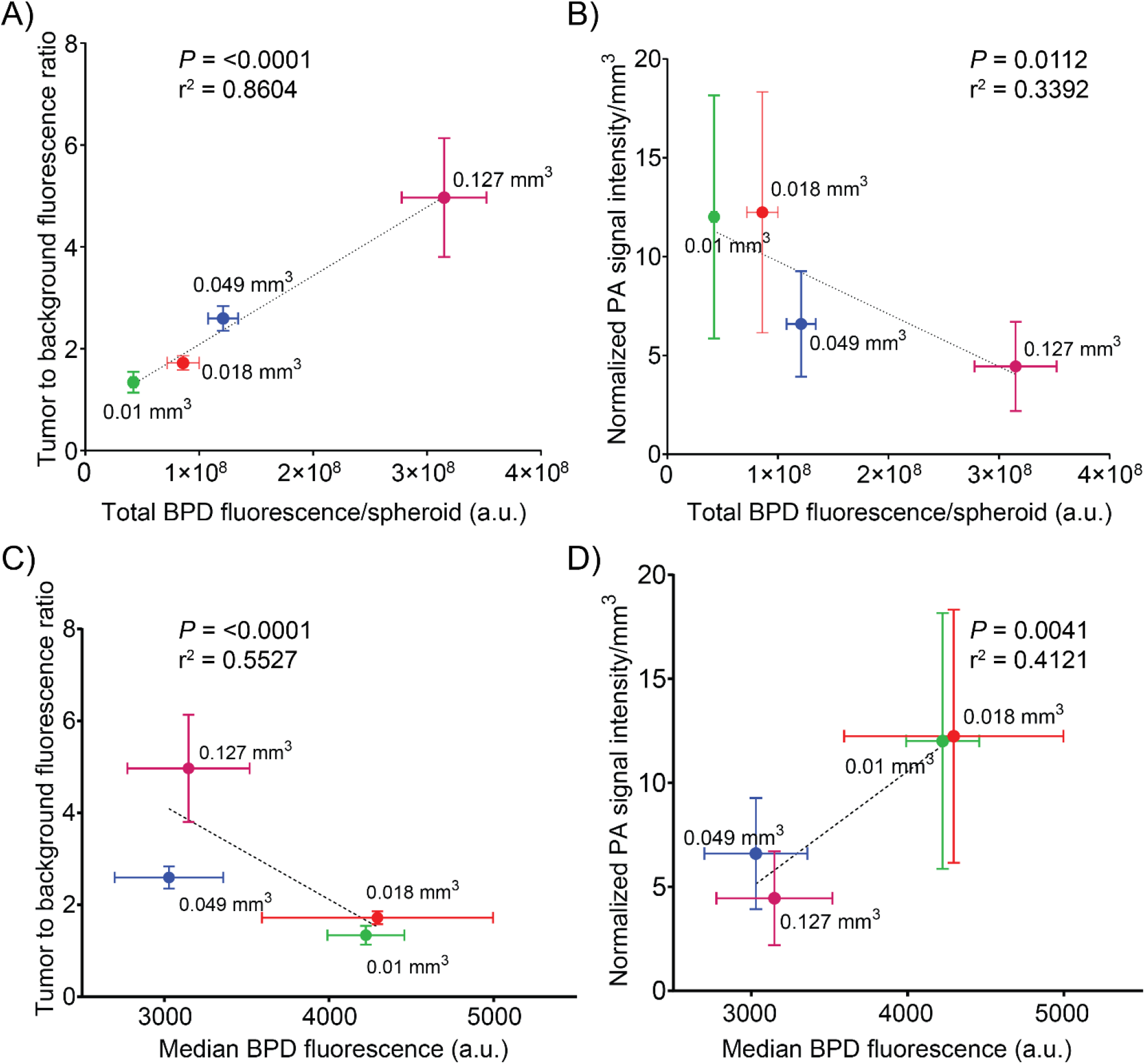
Correlation of tumor spheroid fluorescence to background ratio and photoacoustic signal intensity/mm^3^ with total DFAC (BPD) fluorescence and median BPD fluorescence: Correlations between tumor spheroid fluorescence to background ratio and **A)** Total DFAC (BPD) content, **C)** median BPD fluorescence. While tumor spheroid fluorescence to background ratio positively correlated with total BPD content (*P* < 0.0001, r^2^ = 0.8604), it negatively correlated with median BPD fluorescence (per cell) (*p* < 0.0001, r^2^ = 0.5527). Correlations between photoacoustic signal intensity/mm3 and **B)** Total DFAC (BPD) content, **D)** median BPD fluorescence. While photoacoustic signal intensity/mm^3^ negatively correlated with total DFAC (BPD) content (*p* = 0.0112, r^2^ = 0.3394), it positively correlated with DFAC (median BPD fluorescence) per cell (*p* = 0.0041, r^2^ = 0.4121). Values are mean ± SD; correlations were assessed using Pearson’s r correlation coefficients; n ≥ 3 per condition.

### Dose dependent phototoxicity of tumor spheroids

For studying PDT dose dependent toxicity on the tumor spheroids formed with different cell numbers, the scheme presented in **Figure 6A** was followed. Different spheroids, after a 24 h incubation with DFAC, were irradiated with increasing light doses and longitudinal fluorescence imaging was performed to monitor EGFP fluorescence from the spheroids and calculate viability. As shown in **Figure 6B**, fluorescence intensity of the spheroids decreased with increasing light dose. Quantification of the fluorescence images are provided in **Figures 6C and Supplementary Figure 2A-D**. The decrease in fluorescence intensity was significantly higher for spheroids with lower number of cells, than with spheroids with higher number of cells. At the highest light dose of 100 J/cm^2^, the relative viability of spheroids with smaller volumes (<0.127 mm^3^) was less than 0.1. However, for larger spheroids with volume 0.127 mm^3^, the relative viability was found to be 0.166 ± 0.114. A higher dose was however not tried as these were clinically relevant doses (50 - 100 J/cm^2^ for superficial lesions and 100 - 200 J/cm for interstitial PDT) (NCT02422979, NCT03769506) [4–6, 35]. These results provide insights into the minimal tumor size at which PDT could have a complete response.

**Figure 6:**
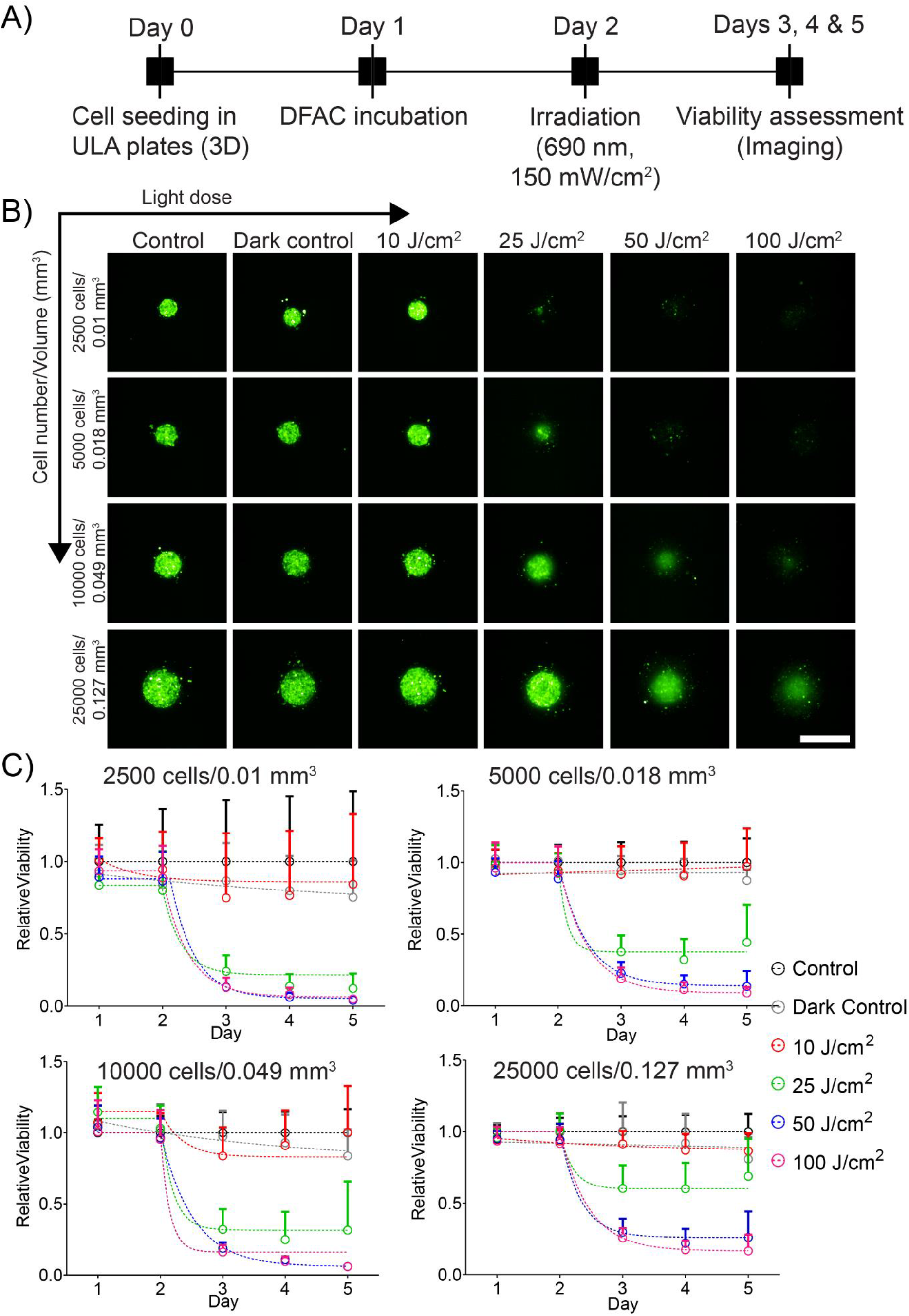
Response of SCC4 spheroids to DFAC-mediated PIT. **A)** Experimental schedule for PIT of SCC4 spheroids. Cells were seeded in CellCarrier Spheroid ULA 96-well microplates followed by incubation with DFAC for 24 h. The spheroids were then irradiated with varying light doses of 10 - 100 J/cm^2^ followed by imaging for 3 days. **B)** Maximum intensity projection (MIP) of fluorescence signals from SCC4 spheroids on day 3 post-PDT. Scale bar corresponds to 500 μm. **C)** Relative viability of SCC4 spheroids treated with PIT. Data are presented as mean ± SD (n ≥ 10 spheroids per condition per time-point).

### Tumor recurrence

The spheroids subjected to PDT were imaged longitudinally for 4-5 weeks and recurrence was determined by quantifying fluorescence intensity of the spheroids at different time-points during the course of the study (**Figure 7A**). **Figure 7B** shows the images of the different spheroids on different days post-treatment with 100 J/cm^2^. As evident, smaller spheroids (0.01 mm^3^ and 0.018 mm^3^) showed no recurrence while larger spheroids (0.049 mm^3^ and 0.127 mm^3^) showed significant recurrence during the course of study. The quantification of relative viabilities of the different spheroids treated with increasing light doses is shown in **Figure 7C and Supplementary Figure 3A-D**. Recurrence was higher for larger spheroids irrespective of the light doses used for treatment, while recurrence was lower for smaller spheroids treated with higher light doses.

**Figure 7:**
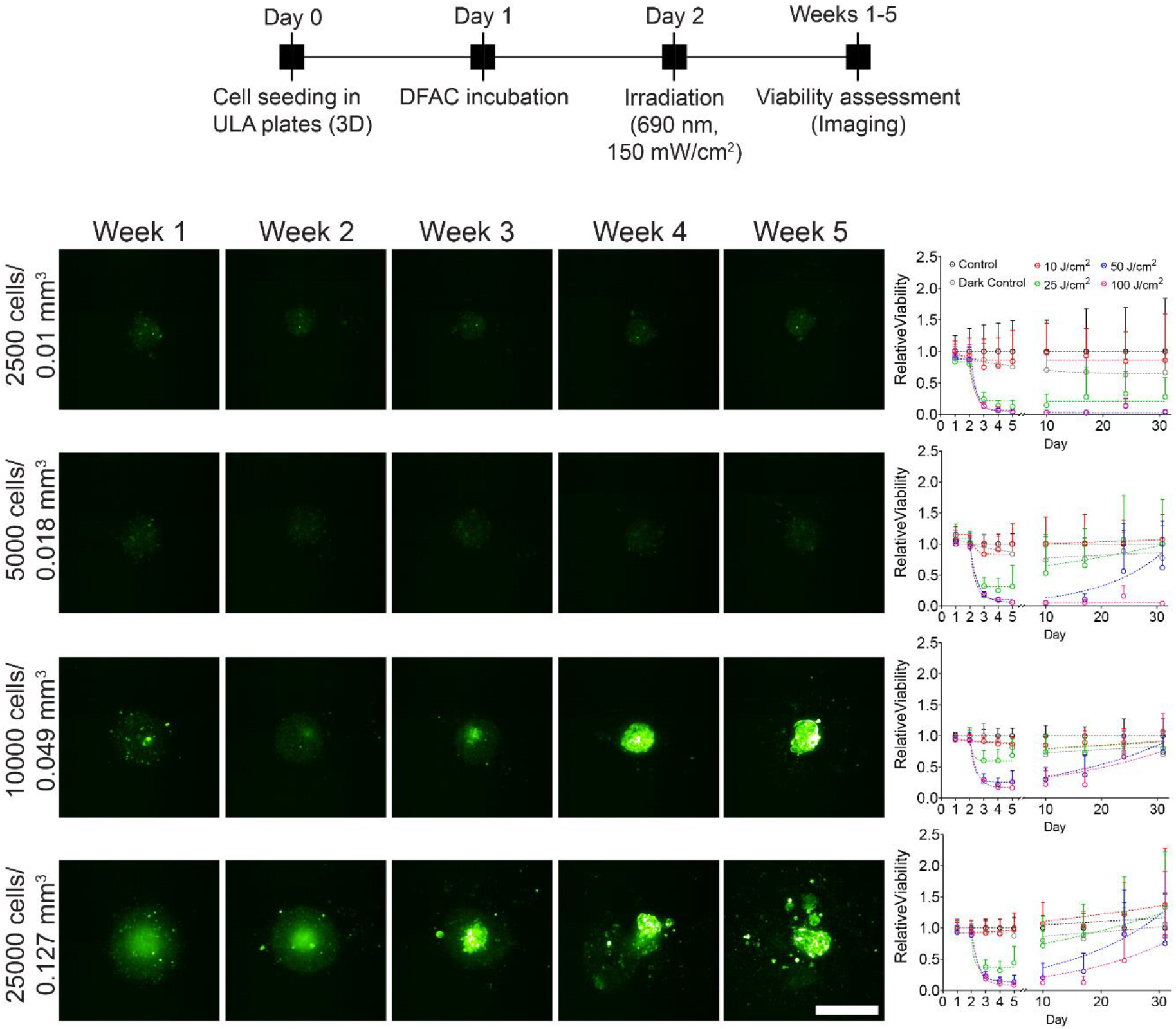
Long-term response of SCC4 spheroids to DFAC-mediated PIT. **A)** Experimental schedule for long-term response of SCC4 spheroids to PIT. Cells were seeded in CellCarrier Spheroid ULA 96-well microplates followed by incubation with DFAC for 24 h. The spheroids were then irradiated with varying light doses of 10 - 100 J/cm^2^ followed by imaging for 5 weeks. **B)** Maximum intensity projection (MIP) of fluorescence signals from SCC4 spheroids on day 31 post-PDT. Scale bar corresponds to 500 μm. Relative viability of SCC4 spheroids treated with PIT are shown in the right panel.

We calculated the recurrence proportion for these spheroids as explained in the methods section. As shown in **figure 8A**, the recurrence proportion for spheroids treated with lower light doses (10 J/cm^2^) was 1, except for the 0.01 mm^3^ spheroids which showed a recurrence proportion of 0.83.

**Figure 8:**
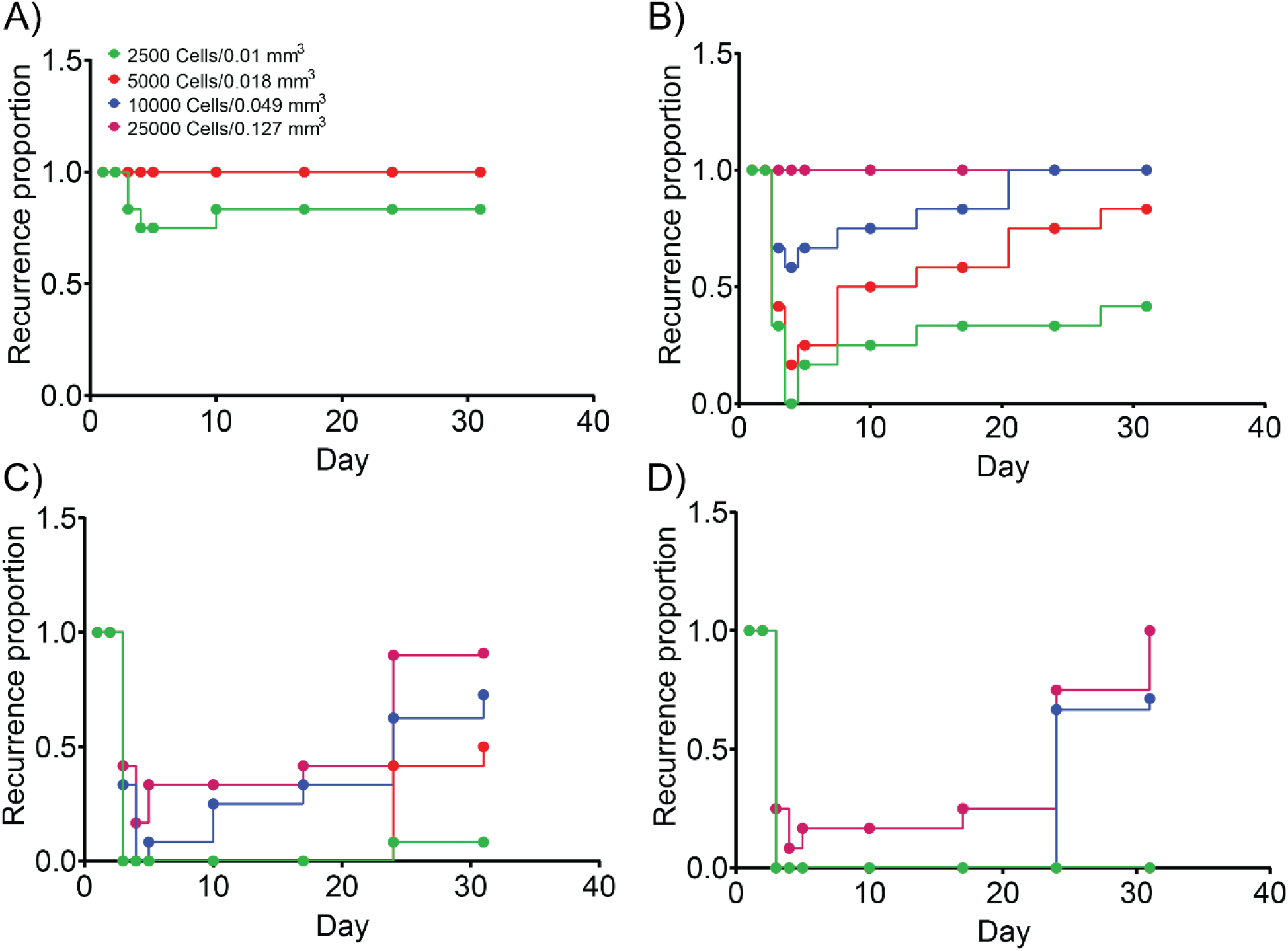
Recurrence of SCC4 spheroid growth after DFAC-mediated PIT. SCC4 spheroids with different volumes were treated with PIT with different light doses **A)**5 J/cm^2^, **B)**25 J/cm^2^, **C)**50 J/cm^2^, and **D)**100 J/cm^2^. Recurrence proportion was calculated by counting the number of spheroids with a relative viability of ≥ 0.3 and dividing it by the total number of spheroids. Spheroids with smaller volumes showed a low recurrence at higher light doses, while the spheroids with higher volumes showed a high recurrence.

With increasing light dose the recurrence proportion decreased (**Figures 8B and 8C**), and for the highest light dose tested (100 J/cm^2^), only the larger spheroids (0.049 mm^3^ and 0.127 mm^3^) showed recurrence that too at a later stage (day 24). By day 31, a recurrence proportion of 0.71 and 1 was observed for the 0.049 mm^3^ and 0.127 mm^3^ spheroids, respectively.

## Discussion

Surgical resection followed by chemo-radiation therapy is the standard treatment regimen for head and neck cancers [14]. Current procedures for surgical resection of tumor tissues relies on tactile cues and visual assessment by the surgeon and are therefore subjective [36–38]. In recent years, the introduction of optical imaging using fluorescent contrast agents has improved visualization of tumor margins in real-time [38]. ICG, MB, and derivatives of 5-ALA which are specifically converted by tumor cells to the fluorescent product protoporphyrin IX (PpIX), have been clinically approved for fluorescence image-guided surgery [39–41]. Several antibody-conjugated fluorophores are also in clinical trials for image-guided surgeries specifically for head and neck cancers (NCT04511078, NCT01987375) [42]. However, signal attenuation due to light scattering by tissues, a general lack of depth profiling by fluorescence imaging and the presence of microscopic tumor tissues, after surgical resection, are challenges that have to be addressed to improve outcomes in head and neck cancer management [43]. We previously, reported the use of a dual function antibody conjugate (DFAC) enabling the combination of fluorescence and photoacoustic molecular imaging in tumor phantoms with an enhanced capability to provide volumetric tumor imaging and the ability to treat residual disease, in a single intraoperative imaging session, by photoimmunotherapy [1]. In this study, we demonstrate the utility of the probe – DFAC, in detecting tumor nodules of sizes as small as 0.01 mm^3^ and photoimmunotherapy-based treatment with low recurrence in a tumor spheroid size-dependent manner. Majority (> 70%) of head and neck cancers are diagnosed late (stage 1 or above) and the tumor size at diagnosis can range from 2 – 4 cm (stage 1 and II) or larger (Stage III and IV) at the time of diagnosis [44]. We envision that DFACs could be useful in clinical settings for enhancing visualization of tumor margins, in all dimensions x, y and z, and provide the much-needed depth profiling that is lacking with fluorescence image-guided surgeries [43]. In this study, we show through a series of in vitro studies, that DFAC can effectively detect small tumor volumes, as low as 0.01 mm^3^ by PA imaging and tumor volumes as small as 0.049 mm^3^ by fluorescence imaging at the concentrations utilized. As the maximum DFAC concertation used in the present study was 100 – 1000-fold lower than the clinically approved dose (250 – 500 mg/m^2^), we expect the DFACs can be used to detect tumor volumes in vivo as well.

Resecting healthy tissue that is beyond 5mm surrounding the tumor tissue is a common practice in current tumor resection surgeries [8]. Importantly, the millimeter and micrometer range sensitivity provided by DFAC-enabled PA and fluorescence imaging is important in delineating tumor margins, where it has been reported that tumor recurrence decreases with every millimeter of healthy tissue, surrounding the tumor tissue, resected [8]. We therefore believe that the submillimeter resolution provided by DFAC can assist in reducing the amount of healthy tissue that is resected thus minimizing the compromise in structure and function that these patients suffer [8]. The tumor volumes estimated through volumetric photoacoustic scans were found to be 2-5-folds higher than the actual tumor nodule size (**Supplementary Figure 4**). This may possibly be due to the artifacts created during the preparation of tumor phantoms. Further studies are however required to evaluate the volumetric resolution provided by DFAC in vivo. Fluorescence imaging due to its superficial imaging capability was not used to calculate tumor spheroid volumes. However, the feasibility of imaging larger tumor volumes through fluorescence is important as current clinical imaging protocols employ sufficiently large working distances for fluorescence imaging enabling the surgeon to illuminate and image larger surgical fields and obtain a so called “birds-eye-view” [10]. In contrast, PA imaging requires the placement of transducers in close proximity to the lesion site to enable viewing of the tumor tissue. Through the probe developed in the present study, we believe the ability to image larger tumor volumes through fluorescence and smaller tumor volumes through PA imaging would fit into the clinical imaging regime of visualizing larger tumors, tumor margins and microscopic diseases with high fidelity.

Apart from enabling visualization of tumor spheroids by photoacoustic and fluorescence imaging, we demonstrate that DFAC can be efficiently used for photoimmunotherapy, inducing cell death in a spheroid size dependent manner. While the potential of PDT, as an adjuvant to surgery, has been established in several previous studies [45–47], the optimal time interval between surgery and PDT is suggested to be approximately 6 weeks [45]. Amongst other factors, such time intervals allow for vascularization of the surgical bed which may assist in the accumulation of the photosensitizer at the target site. Using the same probe for imaging and PDT in a single intraoperative setting holds a distinct advantage in this case. As PDT is affected by depth and oxygenation, the removal of the bulk of tumor through surgical resection before PDT may expose the underlying tumor bed for more effective irradiation. This would also overcome the limitation of hypovascularization and hypoxia that is usually observed in tumor beds (wounds) after surgical excision and may pose limitations for probe distribution and PDT efficacy [45].

With regards to response to PDT, in our study, smaller tumor spheroids appeared to be more responsive with significantly lower recurrence proportions (**Figure 8**), possibly due to a higher PS uptake per cell and a more homogenous PS distribution resulting in a higher PDT dose (**Supplementary Figure 5**). Moreover, smaller spheroids are expected to be better oxygenated as compared to the larger ones, thereby resulting in a higher PDT response [48]. The largest tumor volumes achieved in this in vitro study were 0.149 mm^3^, which is significantly lower than what is expected for microscopic tumors (< 1 cm) in vivo. While DFAC enabled visualization of these microscopic tumor spheroids, their treatment through PIT however did not lead to a complete response, specifically at low light doses, and most spheroids showed recurrence over a 4-week period of study. This is undesired; however, it is to be noted that the treatment outcome of PDT may be different in vivo. Indeed, clinical trials in head and neck cancers using Foscan and in pancreatic cancers using Verteporfin have shown tumor necrosis in regions as deep as 1 – 1.5 cm [49–51], suggesting that the in vitro PDT efficacy may not accurately reflect the in vivo treatment ability and the clinical feasibility of PDT in larger tumors. An alternative for attaining a complete treatment response could be multiple irradiations that may enhance treatment outcomes, as has been reported for the PIT of pre-clinical tumor models [52].

### Conclusion

In conclusion, this study puts forward the case for a molecular targeted theranostic probe for guiding resection surgeries and treatment of residual tumor through photodynamic therapy (photoimmunotherapy). With recent advances in targeted image-guided surgeries and PIT, specifically in context of head and neck cancer, we believe that the proposed DFAC provide a feasible alternative to provide precision in tumor margin delineation and an opportunity to treat residual microscopic disease through PIT in single intra-operative setting. There are several advantages of the probe proposed in the present study; firstly, it enables multi-modal complementary imaging, as is established in our previous study [1], assisting in accurate delineation of surgical margins. Secondly, the use of PIT through the same probe for treating residual disease, if any. Thirdly, the targeted probe can overcome the non-specific nature of photosensitizer formulations and accumulate deep even in desmoplastic tumors as has been shown in our previous studies on stroma-rich pancreatic cancer spheroids with similar targeted probes [2]. The in vitro capacity of DFAC, as established in the present study, warrants further investigation in pre-clinical tumor models to establish its performance as a theranostic probe for the treatment of head and neck cancers with potential applicability in other EGFR over-expressing tumor tissues.

## Acknowledgements

This work was supported by grants from National Institute of Health R01 CA266701 to SM, and P01 CA084203, R01 CA231606, R01 CA226855, R21 CA220143 and S10OD012326 to TH. The authors would like to acknowledge the help from Scott Selfridge and Robert Pawle (Akita Innovations, LLC, a provider of custom designed bioimaging dyes) for providing the SiNC dye.

## Conflict of interest

The authors declare that they have no conflict of interest

## Author Contributions

Conceptualization, M.A.S., S.M., and T.H.; methodology, M.A.S., S.G.G., A.S., and S.M.; formal analysis, M.A.S., S.G.G., A.S., and S.M.; resources, S.M., and T.H; data curation, M.A.S., and S.G.G.; writing—original draft preparation, M.A.S., and S.M.; writing—review and editing, M.A.S., S.G.G., A.S., S.M., and T.H.; supervision, T.H.; project administration, T.H.; funding acquisition, T.H., and S.M.

## Notes

### Competing Interest Statement

The authors have declared no competing interest.

