## Supplementary figures for "A Dual Function Antibody Conjugate Enabled Photoimmunotherapy Complements Fluorescence and Photoacoustic Imaging of Head and Neck Cancer Spheroids"

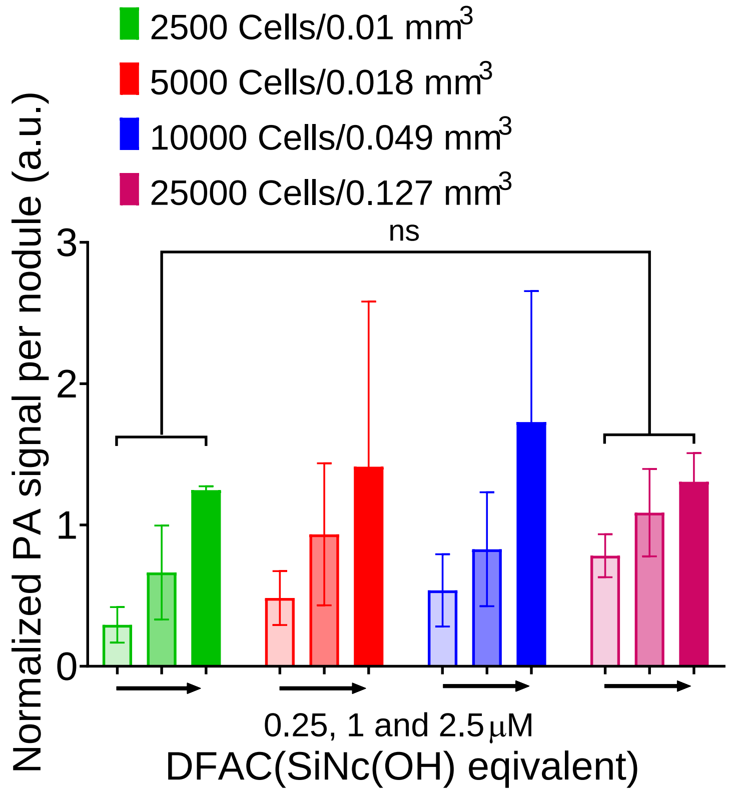


**Supplementary figure 1:** Quantification of photoacoustic signals from the images represented in figure 4A as normalized photoacoustic signal intensity per nodule. Photoacoustic signal from spheroids increased with increase in DFAC concentration, however for spheroids with different sizes treated with the same concentration of DFAC, the normalized photoacoustic signal intensity per nodule remained same. Data are presented as mean ± SD ($n \geq 3$), analyzed using one-way Brown-Forsythe and Welch ANOVA test with post hoc analysis. p-values < 0.05 were considered to be significant and are indicated by asterisks as follows: $^{ns}p > 0.05$, *$p < 0.05$, **$p < 0.01$, ***$p < 0.001$ and ****$p < 0.0001$.

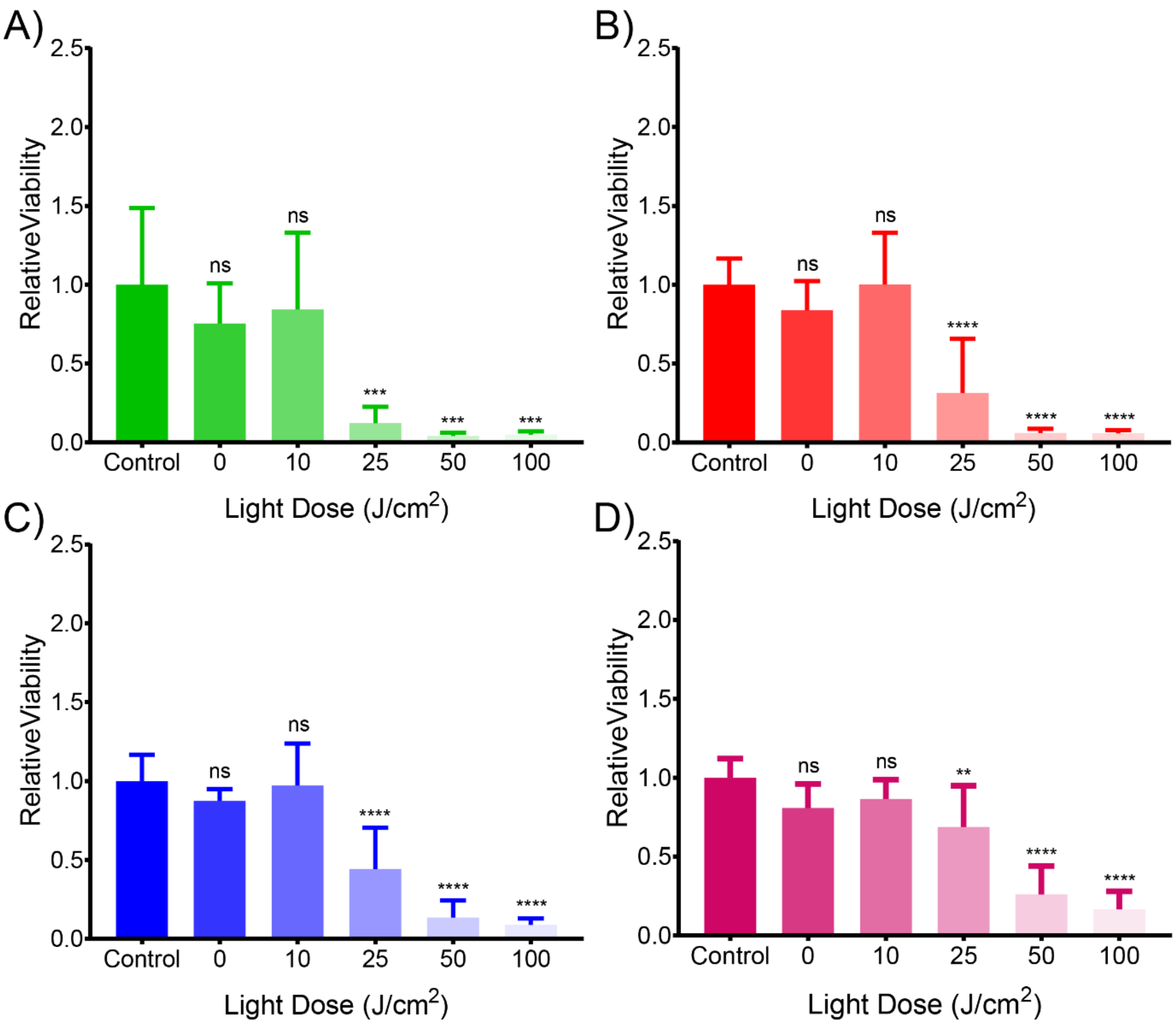


**Supplementary figure 2: Short-term response of SCC4 spheroids to DFAC-mediated PIT.** Relative viability of SCC4 spheroids formed with 2500 **(A)**, 5000 **(B)**, 10000 **(C)**, and 25000 **(D)** cells on day 5 post-PIT. Data are presented as mean ± SD ($n \geq 10$ spheroids per condition per time-point), analyzed using one-way Brown-Forsythe and Welch ANOVA test with post hoc analysis. p-values < 0.05 were considered to be significant and are indicated by asterisks as follows: $^{ns}p > 0.05$, *$p < 0.05$, **$p < 0.01$, ***$p < 0.001$ and ****$p < 0.0001$.

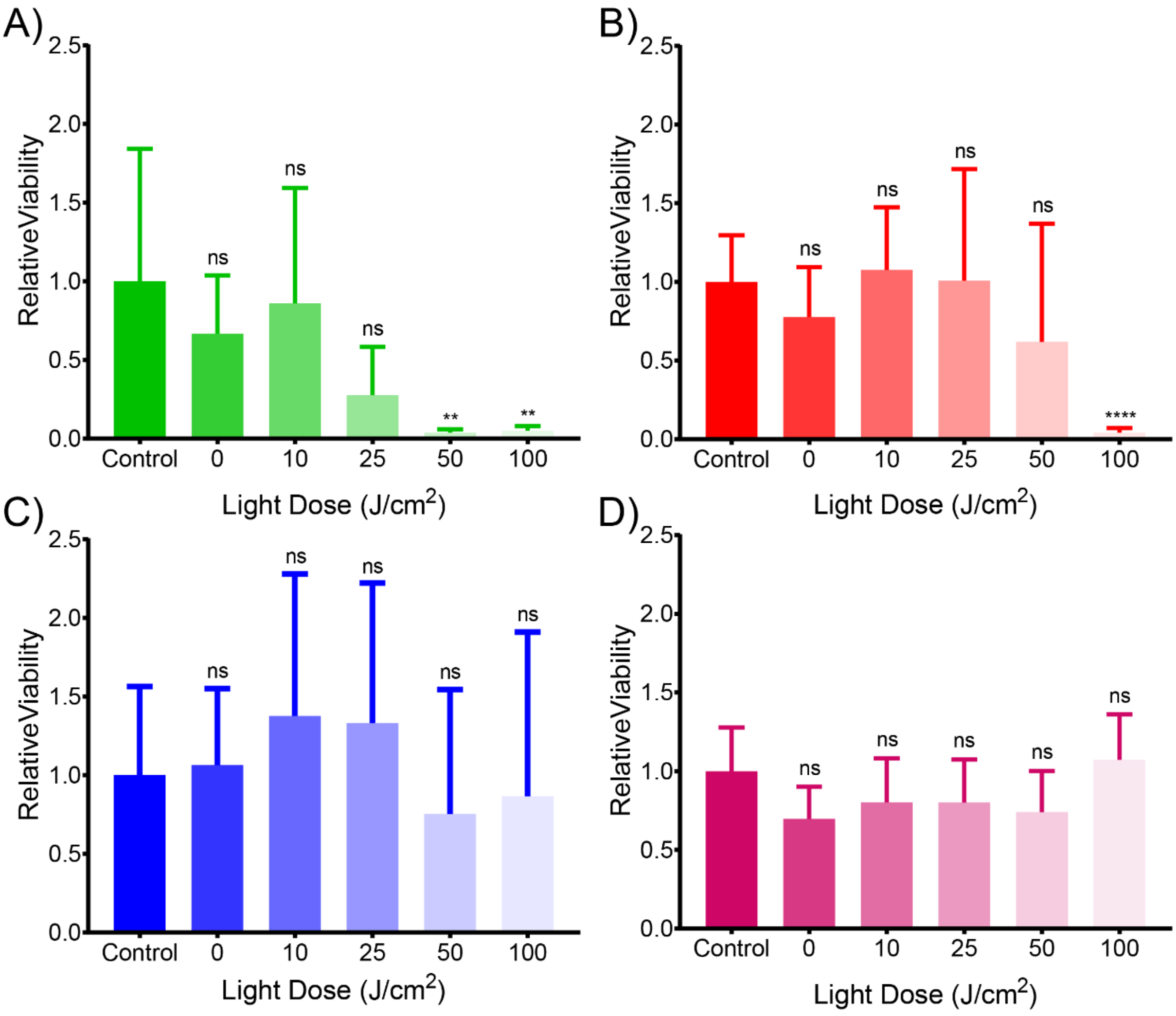


**Supplementary figure 3: Long-term response of SCC4 spheroids to DFAC-mediated PIT.** Relative viability of SCC4 spheroids formed with 2500 **(A)**, 5000 **(B)**, 10000 **(C)**, and 25000 **(D)** cells on day 31 post-PIT. Data are presented as mean ± SD (n ≥ 10 spheroids per condition per time-point), analyzed using one-way Brown-Forsythe and Welch ANOVA test with post hoc analysis. p-values < 0.05 were considered to be significant and are indicated by asterisks as follows: $^{ns}p > 0.05$, *p < 0.05, **p < 0.01, ***p < 0.001 and ****p < 0.0001.

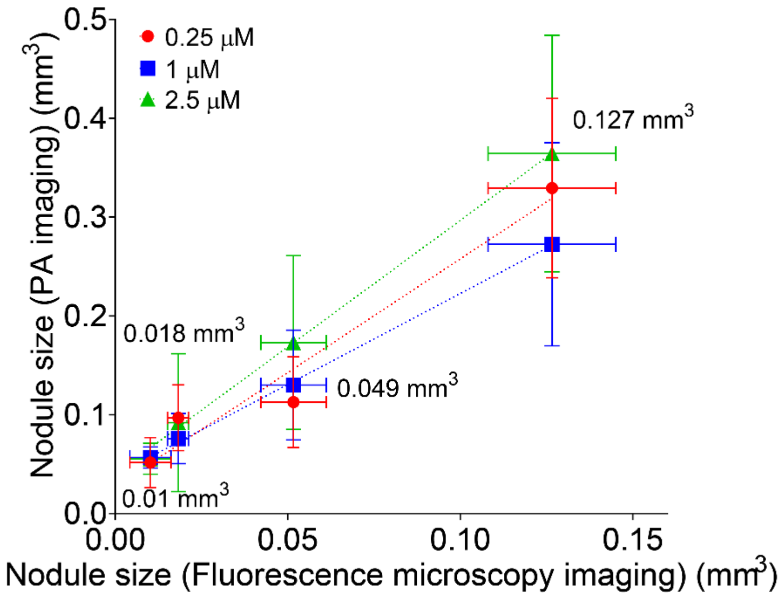


**Supplementary figure 4: Comparison of spheroid sizes as obtained by photoacoustic imaging and fluorescence microscopy imaging.** Photoacoustic imaging consistently over-estimated the size of the 3D spheroids possibly due to the artifacts created during the preparation of tumor spheroid phantoms.

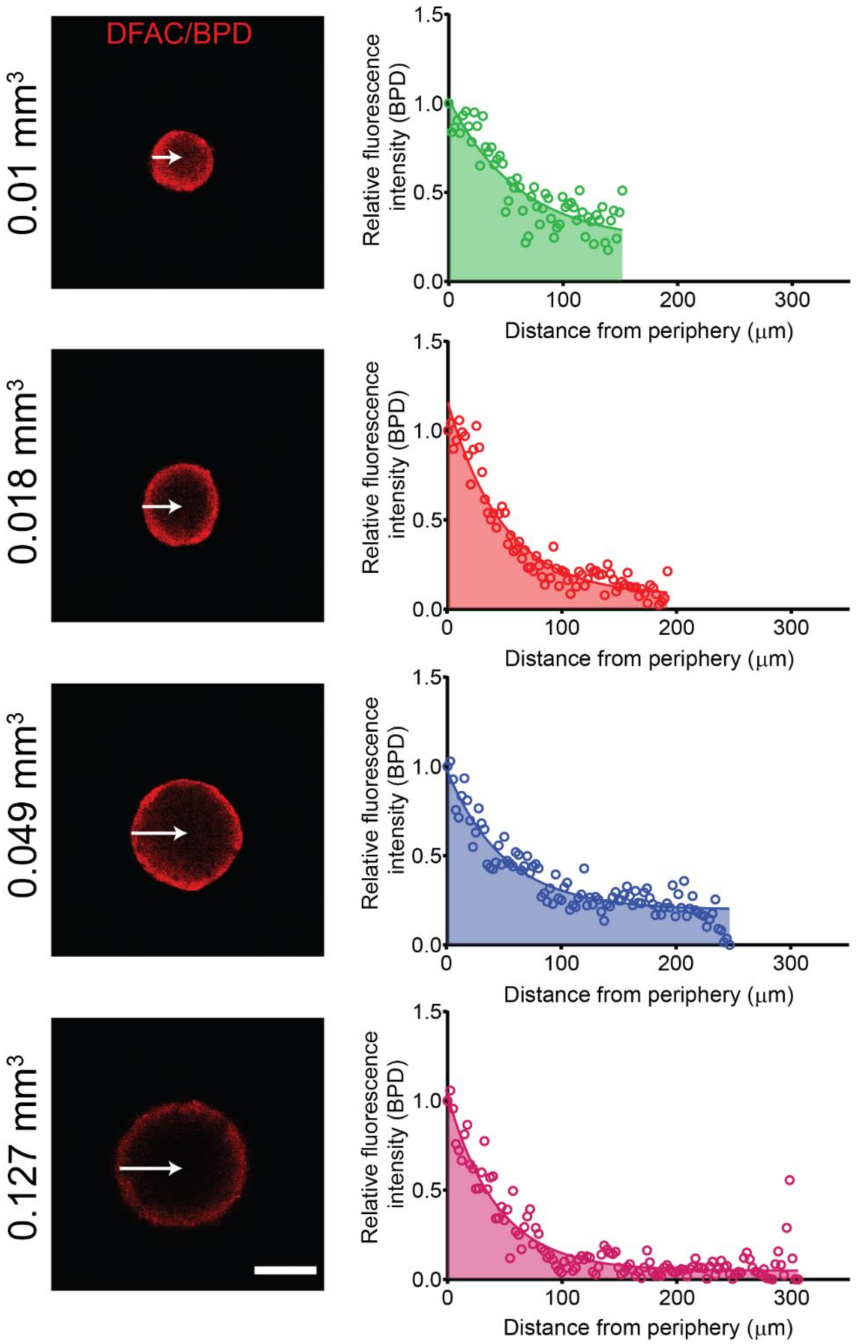


**Supplementary figure 5: Photosensitizer distribution in SCC4 spheroids treated with DFACs:** Representative confocal images of the central optical section of the 3D spheroids. BPD is pseudo-colored as red. Profile plots were generated using image J to monitor BPD distribution across the central plane. As suggested by the profile-plots, BPD concentration appeared to be higher in the spheroids periphery and decreased towards the center of the spheroids. This difference was more pronounced for the larger spheroids which showed an approximately 90% drop in BPD intensity as compared to the smaller spheroids which showed an approximately 50% drop in BPD intensity in the center of the spheroid as compared to the periphery. (Scale bar = 500 μm).
